# The novel coronavirus 2019 (2019-nCoV) uses the SARS-coronavirus receptor ACE2 and the cellular protease TMPRSS2 for entry into target cells

**DOI:** 10.1101/2020.01.31.929042

**Authors:** Markus Hoffmann, Hannah Kleine-Weber, Nadine Krüger, Marcel Müller, Christian Drosten, Stefan Pöhlmann

**Affiliations:** Infection Biology Unit, German Primate Center – Leibniz Institute for Primate Research, Göttingen, Germany; Faculty of Biology and Psychology, University Göttingen, Göttingen, Germany; Institute of Virology, University of Veterinary Medicine Hannover, Hannover, Germany; Research Center for Emerging Infections and Zoonoses, University of Veterinary Medicine Hannover, Hannover, Germany; Charité-Universitätsmedizin Berlin, corporate member of Freie Universität Berlin, Humboldt-Universität zu Berlin, and Berlin Institute of Health, Institute of Virology, Berlin, Germany; German Centre for Infection Research, associated partner Charité, Berlin, Germany; Martsinovsky Institute of Medical Parasitology, Tropical and Vector Borne Diseases, Sechenov University, Moscow, Russia

## Abstract

The emergence of a novel, highly pathogenic coronavirus, 2019-nCoV, in China, and its rapid national and international spread pose a global health emergency. Coronaviruses use their spike proteins to select and enter target cells and insights into nCoV-2019 spike (S)-driven entry might facilitate assessment of pandemic potential and reveal therapeutic targets. Here, we demonstrate that 2019-nCoV-S uses the SARS-coronavirus receptor, ACE2, for entry and the cellular protease TMPRSS2 for 2019-nCoV-S priming. A TMPRSS2 inhibitor blocked entry and might constitute a treatment option. Finally, we show that the serum form a convalescent SARS patient neutralized 2019-nCoV-S-driven entry. Our results reveal important commonalities between 2019-nCoV and SARS-coronavirus infection, which might translate into similar transmissibility and disease pathogenesis. Moreover, they identify a target for antiviral intervention.

**One sentence summary:** The novel 2019 coronavirus and the SARS-coronavirus share central biological properties which can guide risk assessment and intervention.

---

Several members of the family *Coronaviridae* constantly circulate in the human population and usually cause mild respiratory disease (*1*). In contrast, the severe acute respiratory syndrome-associated coronavirus (SARS-CoV) and the Middle East respiratory syndrome-associated coronavirus (MERS-CoV) are transmitted from animals to humans and cause severe respiratory diseases in afflicted human patients, SARS and MERS, respectively (*2*). SARS emerged in 2002 in Guangdong province, China, and its subsequent global spread was associated with 8096 cases and 774 deaths (*3, 4*). The virus uses Chinese horseshoe bats as natural reservoir (*5, 6*) and is transmitted via intermediate hosts to humans. Thus, SARS-CoV was identified in Civet cats and raccoon dogs, which are sold as food sources in Chinese wet markets (*7*). No specific antivirals or approved vaccines are available to combat SARS and the SARS pandemic in 2002/2003 was stopped by conventional control measures, including travel restrictions and patient isolation.

In December 2019 a new infectious respiratory disease emerged in Wuhan, Hubei province, China (*8-10*). Initial infections occurred at Huanan seafood market, potentially due to animal contact. Subsequently, human-to-human transmission occurred (*11*) and the disease rapidly spread within China. A novel coronavirus, 2019-nCoV, which is closely related to SARS-CoV, was detected in patients and is believed to be the etiologic agent of the new lung disease (*10*). On January 28, 2020, at total of 4593 laboratory confirmed infections were reported, including 976 severe cases and 106 deaths (*12*). Infections were also detected in 14 countries outside China and were associated with international travel. At present, it is unknown whether the sequence similarities between 2019-nCoV and SARS-CoV translate into similar biological properties, including pandemic potential (*13*).

The spike (S) protein of coronaviruses facilitates viral entry into target cells. Entry depends on S protein binding to a cellular receptor and on S protein priming by a cellular protease. SARS-S engages angiotensin-converting enzyme 2 (ACE2) as entry receptor (*14*) and employs the cellular serine protease TMPRSS2 for S protein priming (*15-17*). The SARS-S/ACE2 interface has been elucidated and the efficiency of ACE2 usage was found to be a key determinant of SARS-CoV transmissibility (*6, 18*). SARS-S und 2019-nCoV-S share ~76% amino acid identity. However, it is unknown whether 2019-nCoV-S like SARS-S employs ACE2 and TMPRSS2 for host cell entry.

Replication-defective vesicular stomatitis virus (VSV) particles bearing coronavirus S proteins faithfully reflect key aspects of coronavirus host cell entry (*19*). We employed VSV pseudotypes bearing 2019-nCoV-S to study cell entry of 2019-nCoV. Both 2019-nCoV-S and SARS-S were comparably expressed (Fig. 1A) and incorporated into VSV particles (Fig. 1B), allowing a meaningful side-by-side comparison. We first focused on 2019-nCoV cell tropism. Transduction of cell lines of animal and human origin revealed that all cell lines were readily susceptible to entry driven by the pantropic VSV glycoprotein (VSV-G) (Fig. 1C), as expected. (Fig. 1C). Notably, 2019-nCoV-S facilitated entry into an identical spectrum of cell lines as SARS-S (Fig. 1C), suggesting similarities in receptor choice.

**Fig. 1.**
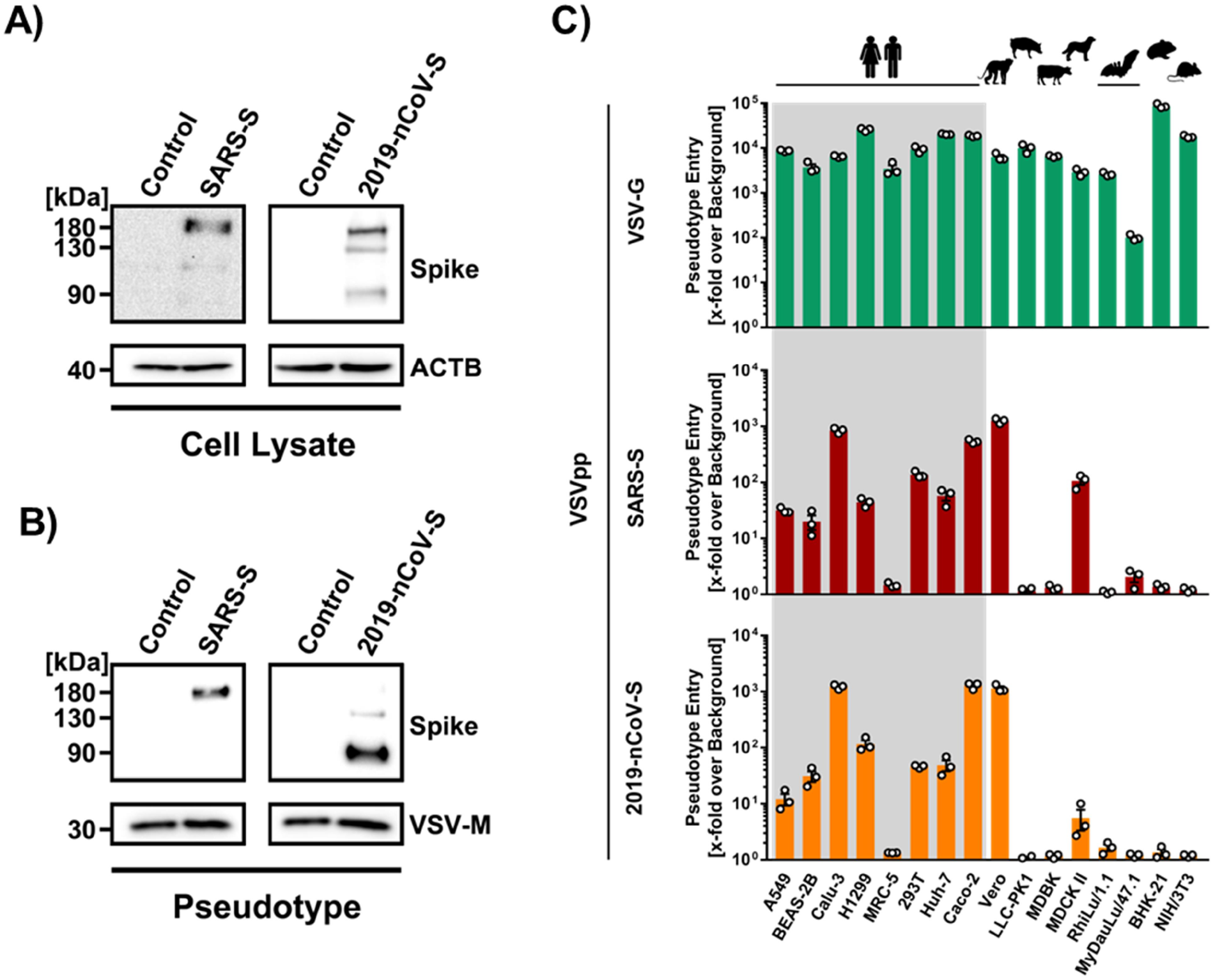
2019-nCoV-S and SARS-S facilitates entry into a similar panel of mammalian cell lines. Analysis of 2019-nCoV-S expression (**A**) and pseudotype incorporation (**B**) by Western blot. Representative blots from three experiments are shown. ß-Actin (cell lysates) and VSV-M (particles) served as loading controls. (C) Cell lines of human and animal origin were inoculated with pseudotyped VSV harboring VSV-G, SARS-S or 2019-nCoV-S. At 16 h postinoculation, pseudotype entry was analyzed. Shown are the combined data of three experiments. Error bars indicate SEM.

Sequence analysis revealed that 2019-nCoV clusters with SARS-CoV-related viruses from bats (SARSr-CoV), of which some but not all can use ACE2 for host cell entry (Fig. 2A and fig. S1). Analysis of the receptor binding motif (RBM), a portion of the receptor binding domain (RBD) that makes contact with ACE2, revealed that most amino acid residues essential for ACE2 binding were conserved in 2019-nCoV-S but not in the S proteins of SARSr-CoV previously found not to use ACE2 for entry (Fig. 2B). In agreement with these findings, directed expression of human and bat ACE2 but not human DPP4, the entry receptor used by MERS-CoV (*20*), or human APN, the entry receptor used by HCoV-229E (*21*), allowed 2019-nCoV-S- and SARS-S-driven entry into otherwise non-susceptible BHK-21 cells (Fig. 2C), indicating that 2019-nCoV-S like SARS-S uses ACE2 for cellular entry.

**Fig. 2.**
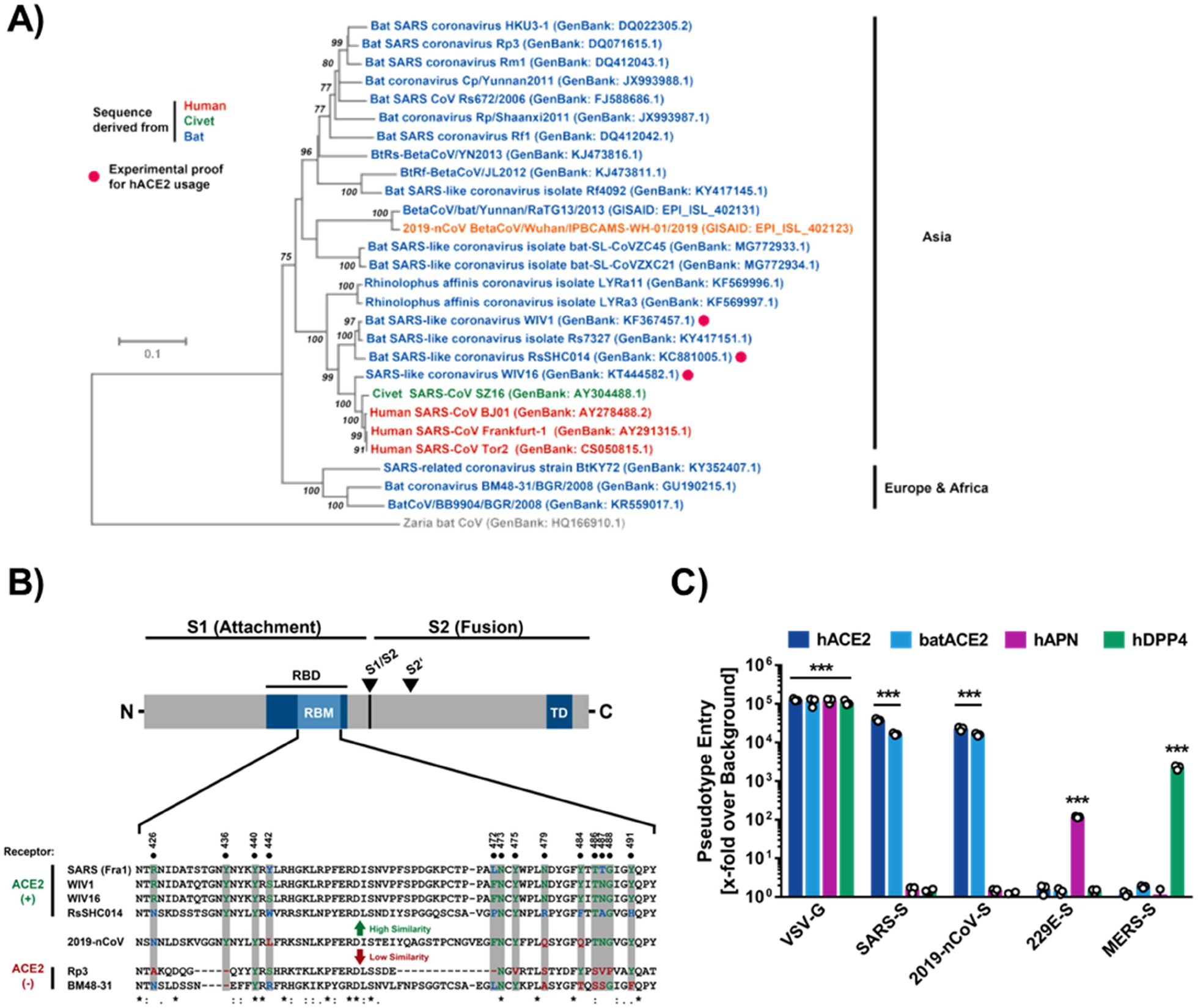
2019-nCoV-S utilizes ACE2 as cellular receptor. (**A**) The S protein of 2019-nCoV clusters phylogenetically with S proteins of known bat-associated betacoronaviruses (see also SI Figure 1 for more details). (**B**) Alignment of the receptor binding motif of SARS-S with corresponding sequences of bat-associated betacoronavirus S proteins that are able or unable to use ACE2 as cellular receptor reveals that 2019-nCoV possesses amino acid residues crucial for ACE2 binding. (**C**) 293T cells transiently expressing ACE2 of human (dark blue) or bat (light blue) origin, human APN (purple) or hDPP4 (green) were inoculated with pseudotyped VSV harboring VSV-G, SARS-S, 2019-nCoV-S, MERS-S or 229E-S. At 16 h postinoculation, pseudotype entry was analyzed. The average of three independent experiments is shown. Error bars indicate SEM.

We next investigated protease dependence of 2019-nCoV entry. SARS-CoV can use the endosomal cysteine proteases cathepsin B and L (CatB/L) for S protein priming in TMPRSS2^-^ cell lines (*22*). However, TMPRSS2 is expressed in viral target cells in the lung (*23*) and entry into TMPRSS2^+^ cell lines is promoted by TMPRSS2 (*15-17*) and is partially CatB/L independent, although blockade of both proteases is required for efficient entry inhibition (*24*). Moreover, TMPRSS2 but not CatB/L activity is essential for spread of SARS-CoV and other coronaviruses in the infected host (*25, 26*). For initial insights into 2019-nCoV-S protease choice, we employed ammonium chloride, which elevates endosomal pH and thereby blocks CatB/L activity. Ammonium chloride treatment blocked VSV-G-driven entry into both cell lines studied while entry driven by the Nipah virus F and G proteins was not affected (Fig. 3A), in keeping with expectations. Moreover, ammonium chloride treatment strongly inhibited 2019-nCoV-S- and SARS-S-driven entry into TMPRSS2^-^ 293T cells while inhibition of entry into TMPRSS2^+^ Caco-2 cells was less efficient, which would be compatible with 2019-nCoV-S priming by TMPRSS2 in Caco-2 cells. Indeed, the serine protease inhibitor camostat mesylate, which is active against TMPRSS2 (*24*), efficiently blocked 2019-nCoV-S-driven entry into Caco-2 (TMPRSS2^+^) but not 293T (TMPRSS2^-^) cells while the CatB/L inhibitor E64d had the opposite effect (Fig. 3B). Moreover, directed expression of TMPRSS2 rescued 2019-nCoV-S-driven entry from inhibition by E64d (Fig. 3C), demonstrating that 2019-nCoV-S uses TMPRSS2 for priming.

**Fig. 3.**
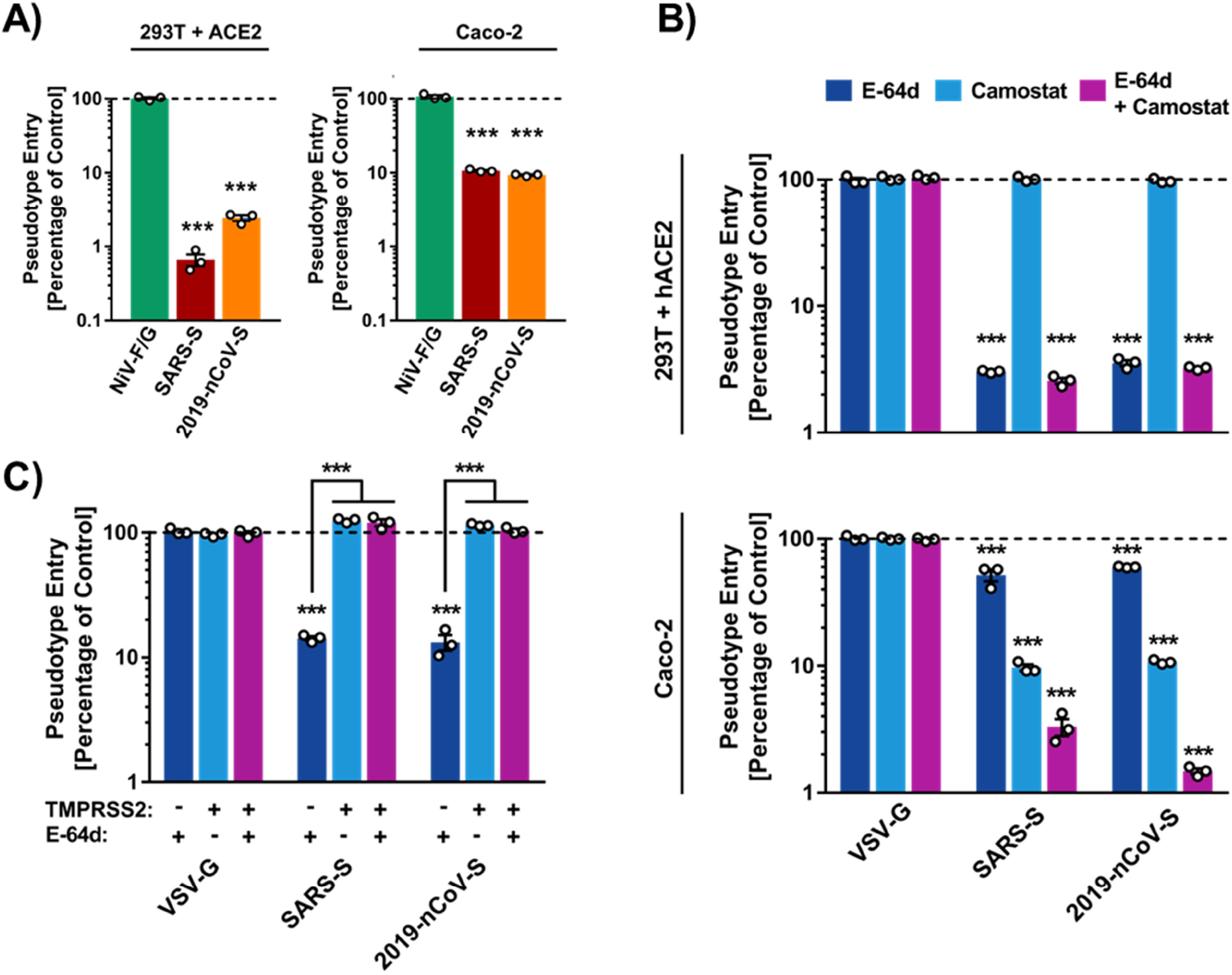
2019-nCoV-S employs TMPRSS2 for S protein priming. Ammonium chloride (**A**), E64d (CatB/L inhibitor) (**B**) and/or camostat (TMPRSS2 inhibitor) (**B**) were added to the indicated target cells before transduction with pseudotypes bearing the indicated glycoproteins. (**C**) 293T cells transiently expressing ACE2 alone or in combination with TMPRSS2 were incubated with CatB/L inhibitor E64d or PBS as control and inoculated with pseudotypes bearing the indicated viral surface proteins. The average of three independent experiments is shown in panels A-C. Error bars indicate SEM. Statistical significance was tested by two-way ANOVA with Dunnett posttest.

Convalescent SARS patients exhibit a neutralizing antibody response directed against the viral S protein (*27*). We investigated whether such antibodies block 2019-nCoV-S-driven entry. Serum from a convalescent SARS patient inhibited SARS-S-but not VSV-G-driven entry in a concentration dependent manner (Fig. 4). In addition, the serum reduced 2019-nCoV-S-driven entry, although with somewhat lower efficiency as compared to SARS-S (Fig. 4). Thus, antibody responses raised against SARS-S during infection or vaccination might offer some protection against 2019-nCoV infection.

**Fig. 4.**
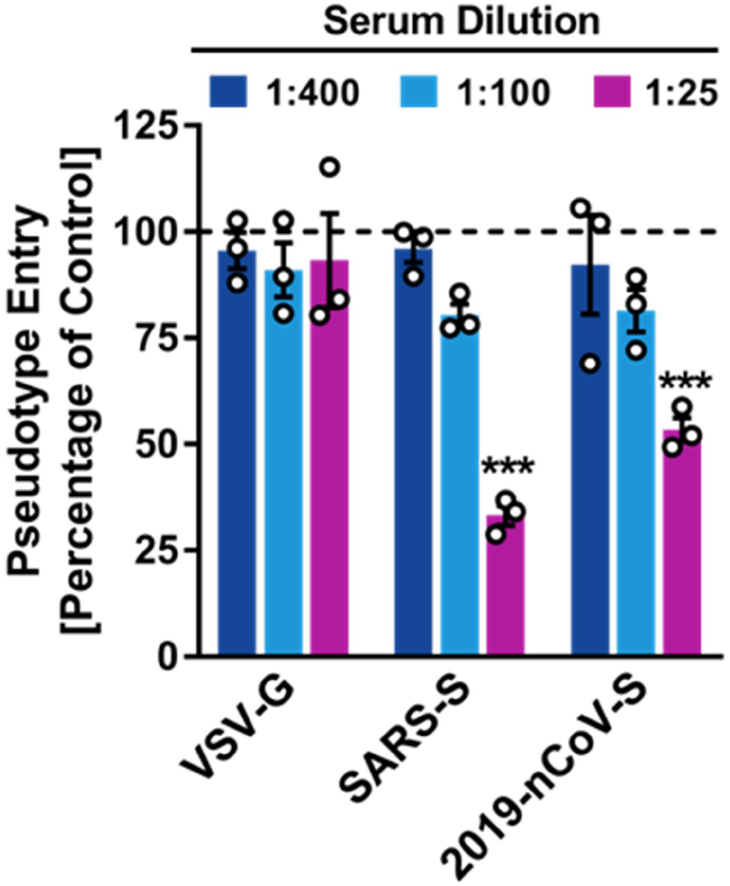
Serum from a convalescent SARS patient cross-neutralizes 2019-nCoV-S-driven entry. Pseudotypes harboring the indicated viral surface proteins were incubated with different dilutions of serum from a convalescent SARS patient and subsequently inoculated onto 293T cells that transiently express ACE2 in order to evaluate cross-neutralization. The results from a representative experiment with triplicate samples are shown and were confirmed in a separate experiment. Error bars indicate SD. Statistical significance was tested by two-way ANOVA with Dunnett posttest.

The finding that 2019-nCoV-S and SARS-S use the same receptor, ACE2, for entry into target cells has important implications for our understanding of 2019-nCoV transmissibility and pathogenesis. Thus, one can expect that 2019-nCoV targets the same cells as SARS-CoV and that the previously documented modest ACE2 expression in the upper respiratory tract (*23, 28*) might limit 2019-nCoV transmissibility. Moreover, it is noteworthy that ACE2 expression is not limited to the lung and that extrapulmonary spread of SARS-CoV in ACE2^+^ tissues was observed (*29, 30*). The same can be expected for 2019-nCoV, although affinity of SARS-S and 2019-nCoV-S for ACE2 remains to be compared.

Priming of coronavirus S proteins by host cell proteases is essential for viral entry into cells and protease choice can determine zoonotic potential (*31*). The S proteins of SARS-CoV can use the endosomal cysteine proteases for S protein priming in TMPRSS2^-^ cells (*22*). However, S protein priming by TMPRSS2 but not CatB/L is essential for viral entry into primary target cells and for viral spread in the infected host (*24-26*). The present study suggests that 2019-nCoV spread might also depend on TMPRSS2 activity and it is noteworthy that the serine protease inhibitor camostat mesylate blocks TMPRSS2 activity (*24, 26*) and has been approved in Japan for human use, although for an unrelated indication. This compound or related ones should be considered for treatment of 2019-nCoV infected patients.

Convalescent SARS patients exhibit a neutralizing antibody response that can be detected even 24 months after infection (*27*) and this is largely directed against the S protein. Moreover, experimental SARS vaccines, including recombinant S protein (*32*) and inactivated virus (*33*) induce neutralizing antibody responses. Our results, although limited in significance due to a single patient serum being available for testing, indicate that neutralizing antibody responses raised against SARS-S should also offer some protection against 2019-nCoV infection, which may have implications for outbreak control.

## Acknowledgments

We thank Heike Hofmann-Winkler for advice.

## Funding

This work was supported by BMBF (RAPID Consortium, 01KI1723D and 01KI1723A).

## Authors contributions

M.H. and S.P. designed the study. S.P. and C.D. acquired funding for the study. M.H. and H.K.-W. conducted experiments and analyzed data. N.K., M.M. and C.D. provided important resources. M.H. and S.P. wrote the manuscript with input from all authors.

### Competing interests

The authors declare no competing interests.

### Data and materials availability

All materials used in this study will be provided upon signature of an appropriate material transfer agreement. All data are available in the main text or the supplementary materials

## Supplementary Materials

### Cell Culture

All cell lines were incubated at 37 °C and 5 % CO_2_ in a humidified atmosphere. 293T (human, kidney), BHK-21 (Syrian hamster, kidney cells), Huh-7 (human, liver), LLC-PK1 (pig, kidney), MRC-5 (human, lung), MyDauLu/47.1 (Daubenton’s bat *[Myotis daubentonii]*, lung), NIH/3T3 (Mouse, embryo), RhiLu/1.1 (Halcyon horseshoe bat *[Rhinolophus alcyone]*, lung), Vero (African green monkey, kidney) cells were incubated in Dulbecco’s’ modified Eagle medium (PAN-Biotech). Calu-3 (human, lung), Caco-2 (human, colon), MDBK (cattle, kidney) and MDCK II (Dog, kidney) cells were incubated in Minimum Essential Medium (ThermoFisher Scientific). A549 (human, lung), BEAS-2B (human, bronchus) and NCI-H1299 (human, lung) cells were incubated in DMEM/F-12 Medium with Nutrient Mix (ThermoFisher Scientific). All media were supplemented with 10 % fetal bovine serum (Biochrom), 100 U/ml of penicillin and 0.1 mg/ml of streptomycin (PAN-Biotech), 1x non-essential amino acid solution (10x stock, PAA) and 10 mM sodium pyruvate solution (ThermoFisher Scientific). For seeding and subcultivation, cells were first washed with phosphate buffered saline (PBS) and then incubated in the presence of trypsin/EDTA solution (PAN-Biotech) until cells detached. Transfection was carried out by calcium-phosphate precipitation.

### Plasmids

Expression plasmids for vesicular stomatitis virus (VSV, serotype Indiana) glycoprotein (VSV-G), SARS-S (derived from the Frankfurt-1 isolate) with or without a C-terminal HA epitope tag, HCoV-229E-S, MERS-S, human and bat angiotensin converting enzyme 2, human aminopeptidase N, human dipeptidyl-peptidase 4 and human TMPRSS2 have been described elsewhere (*1-6*). For generation of the expression plasmids for 2019-nCoV-S with or without a C-terminal HA epitope tag we PCR-amplified the coding sequence of a synthetic, codon-optimized (for human cells) 2019-nCoV-S DNA (GeneArt Gene Synthesis, ThermoFisher Scientific) based on the publicly available protein sequence in the National Center for Biotechnology Information database (NCBI Reference Sequence: YP_009724390.1) and cloned in into the pCG1 expression vector via BamHI and XbaI restriction sites.

### Pseudotyping of VSV and transduction experiments

*Pseudotyping*: VSV pseudotypes were generated according to a published protocol (*7*). In brief, 293T transfected to express the viral surface glycoprotein under study were inoculated with a replication-deficient VSV vector that contains expression cassettes for eGFP (enhanced green fluorescent protein) and firefly luciferase instead of VSV-G the open reading frame, VSV*ΔG-fLuc (kindly provided by Gert Zimmer, Institute of Virology and Immunology, Mittelhäusern/Switzerland). After an incubation period of 1 h at 37 °C, the inoculum was removed and cells were washed with PBS before medium supplemented with anti-VSV-G antibody (I1, mouse hybridoma supernatant from CRL-2700; ATCC) was added (no antibody was added to cells expressing VSV-G). Pseudotype particles ware harvested 16 h postinoculation, clarified from cellular debris by centrifugation and used for experimentations.

*Transduction of target cells*: Target cells were grown in 96-well plates until they reached 50-75 % confluency before they were inoculated with the respective pseudotype vectors. For experiments addressing the search for the 2019-nCoV receptor cells were transfected with expression plasmids 24 h in advance. For experiments involving ammonium chloride (final concentration 50 mM) and protease inhibitors (E-64d, 25 μM; camostat mesylate, 100 μM), target cells were treated with the respective chemical 2 h in advance. For neutralization experiments, pseudotypes were pre-incubated for 30 min at 37 °C with different serum dilutions. Transduction efficiency was quantified 16 h posttransduction by measuring the activity firefly luciferase in cell lysates.

### Analysis of 2019-nCoV-S expression and particle incorporation by SDS-PAGE and immunoblot

*Preparation of whole cell lysates:* 293T cells were transfected with expression vectors for HA-tagged 2019-nCoV-S or SARS-S, or empty expression vector (negative control). The culture medium was replaced at 16 h posttransfection and the cells were incubated for an additional 24 h. Then, the culture medium was removed and cells were washed once with PBS before 2x SDS-sample buffer (0.03 M Tris-HCl, 10% glycerol, 2% SDS, 0.2% bromophenol blue, 1 mM EDTA) was added and cells were incubated for 10 min at room temperature. Next, the samples were heated for 15 min at 96 °C and subjected to SDS-PAGE and immunoblotting. *Preparation of pseudotype particle lysates*: 1 ml of the respective VSV pseudotype were loaded on a 20 % (w/v) sucrose cushion (volume 40 μl) and subjected to high-speed centrifugation (25.000 g for 120 min at 4 °C). Thereafter, 1 ml of supernatant was removed and the residual volume was mixed with 40 μl of 2x SDS-sample buffer, heated for 15 min at 96 °C and subjected to SDS-PAGE and immunoblotting

After protein transfer, nitrocellulose membranes were blocked in 5 % skim milk (in PBS containing 0.05 % Tween-20, PBS-T) for 1 h at room temperature and then incubated over night at 4 °C with the primary antibody (diluted in PBS-T). Following three washing intervals of 10 min in PBS-T the membranes were incubated for 1 h at room temperature with the secondary antibody (diluted in PBS-T), before the membranes were washed and imaged using an in in house-prepared enhanced chemiluminescent solution (0.1 M Tris-HCl [pH 8.6], 250 μg/ml luminol, 1 mg/ml para-hydroxycoumaric acid, 0.3 % H_2_O_2_) and the ChemoCam imaging system along with the ChemoStar Professional software (Intas Science Imaging Instruments GmbH). The following primary antibodies were used: Mouse anti-HA tag (Sigma-Aldrich, H3663, 1:2,500), mouse anti-ß-actin (Sigma-Aldrich, A5441, 1:2,000), mouse anti-VSV matrix protein (Kerafast, EB0011, 1:2,500). As secondary antibody we used a peroxidase-coupled goat anti-mouse antibody (Dianova, 115-035-003, 1:10000).

### Phylogenetic analysis

Phylogenetic analysis (neighbor-joining trees) was performed using the MEGA7.0.26 software. Reference sequences were obtained from the National Center for Biotechnology Information and GISAID (Global Initiative on Sharing All Influenza Data) databases. Reference numbers are indicated in the figures.

### Statistical analysis

One-way or two-way analysis of variance (ANOVA) with Dunnett’s or Sidaks’ posttest was used to test for statistical significance. Only p values of 0.05 or lower were considered statistically significant (p > 0.05 [ns, not significant], p ≤ 0.05 [*], p ≤ 0.01 [**], p ≤ 0.001 [***]). For all statistical analyses, the GraphPad Prism 7 software package was used (GraphPad Software).

---

**Supplementary figure S1.**
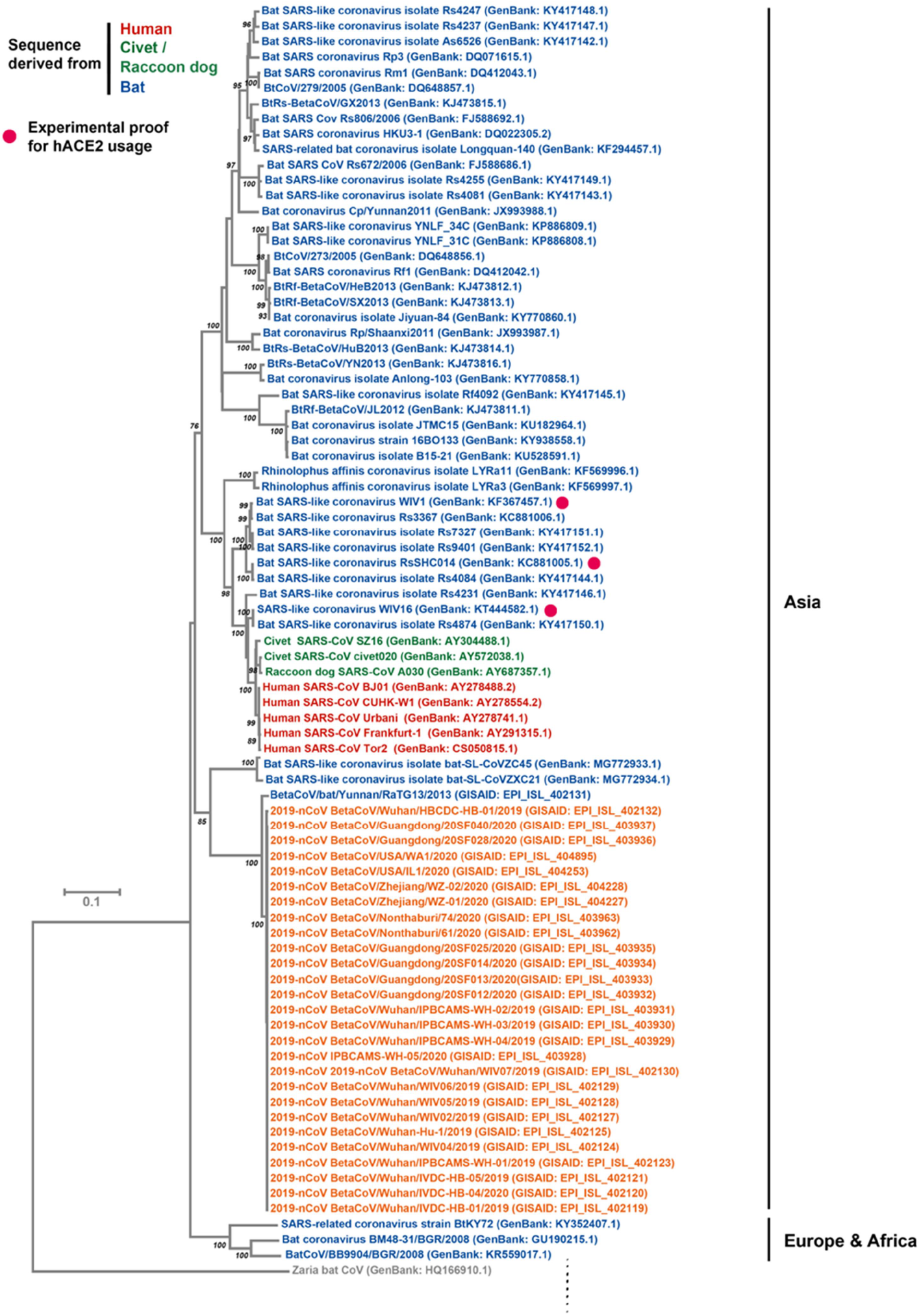
**Legend to supplementary figure S1**. Phylogenetic analysis (neighbor-joining tree) of spike protein sequences. Small numbers indicate bottstrap values (only values higher that 75 are shown).

## References and Notes

1. V. M. Corman, J. Lienau, M. Witzenrath, [Coronaviruses as the cause of respiratory infections]. Internist (Berl) 60, 1136–1145 (2019).

2. A. R. Fehr, R. Channappanavar, S. Perlman, Middle East Respiratory Syndrome: Emergence of a Pathogenic Human Coronavirus. Annu Rev Med 68, 387–399 (2017).

3. W. H. Organization. (2004), vol. 2020.

4. E. de Wit, N. van Doremalen, D. Falzarano, V. J. Munster, SARS and MERS: recent insights into emerging coronaviruses. Nat Rev Microbiol 14, 523–534 (2016).

5. S. K. Lau et al., Severe acute respiratory syndrome coronavirus-like virus in Chinese horseshoe bats. Proc Natl Acad Sci U S A 102, 14040–14045 (2005).

6. F. Li, W. Li, M. Farzan, S. C. Harrison, Structure of SARS coronavirus spike receptor-binding domain complexed with receptor. Science 309, 1864–1868 (2005).

7. Y. Guan et al., Isolation and characterization of viruses related to the SARS coronavirus from animals in southern China. Science 302, 276–278 (2003).

8. C. Wang, P. W. Horby, F. G. Hayden, G. F. Gao, A novel coronavirus outbreak of global health concern. Lancet, (2020).

9. C. Huang et al., Clinical features of patients infected with 2019 novel coronavirus in Wuhan, China. Lancet, (2020).

10. N. Zhu et al., A Novel Coronavirus from Patients with Pneumonia in China, 2019. N Engl J Med, (2020).

11. J. F. Chan et al., A familial cluster of pneumonia associated with the 2019 novel coronavirus indicating person-to-person transmission: a study of a family cluster. Lancet, (2020).

12. W. H. Organization. (2020), vol. 2020.

13. V. J. Munster, M. Koopmans, N. van Doremalen, D. van Riel, E. de Wit, A Novel Coronavirus Emerging in China - Key Questions for Impact Assessment. N Engl J Med, (2020).

14. W. Li et al., Angiotensin-converting enzyme 2 is a functional receptor for the SARS coronavirus. Nature 426, 450–454 (2003).

15. I. Glowacka et al., Evidence that TMPRSS2 activates the severe acute respiratory syndrome coronavirus spike protein for membrane fusion and reduces viral control by the humoral immune response. J Virol 85, 4122–4134 (2011).

16. S. Matsuyama et al., Efficient activation of the severe acute respiratory syndrome coronavirus spike protein by the transmembrane protease TMPRSS2. J Virol 84, 12658–12664 (2010).

17. A. Shulla et al., A transmembrane serine protease is linked to the severe acute respiratory syndrome coronavirus receptor and activates virus entry. J Virol 85, 873–882 (2011).

18. W. Li et al., Receptor and viral determinants of SARS-coronavirus adaptation to human ACE2. EMBO J 24, 1634–1643 (2005).

19. H. Kleine-Weber et al., Mutations in the Spike Protein of Middle East Respiratory Syndrome Coronavirus Transmitted in Korea Increase Resistance to Antibody-Mediated Neutralization. J Virol 93, (2019).

20. V. S. Raj et al., Dipeptidyl peptidase 4 is a functional receptor for the emerging human coronavirus-EMC. Nature 495, 251–254 (2013).

21. C. L. Yeager et al., Human aminopeptidase N is a receptor for human coronavirus 229E. Nature 357, 420–422 (1992).

22. G. Simmons et al., Inhibitors of cathepsin L prevent severe acute respiratory syndrome coronavirus entry. Proc Natl Acad Sci U S A 102, 11876–11881 (2005).

23. S. Bertram et al., Influenza and SARS-coronavirus activating proteases TMPRSS2 and HAT are expressed at multiple sites in human respiratory and gastrointestinal tracts. PLoS One 7, e35876 (2012).

24. M. Kawase, K. Shirato, L. van der Hoek, F. Taguchi, S. Matsuyama, Simultaneous treatment of human bronchial epithelial cells with serine and cysteine protease inhibitors prevents severe acute respiratory syndrome coronavirus entry. J Virol 86, 6537–6545 (2012).

25. N. Iwata-Yoshikawa et al., TMPRSS2 Contributes to Virus Spread and Immunopathology in the Airways of Murine Models after Coronavirus Infection. J Virol 93, (2019).

26. Y. Zhou et al., Protease inhibitors targeting coronavirus and filovirus entry. Antiviral Res 116, 76–84 (2015).

27. W. Liu et al., Two-year prospective study of the humoral immune response of patients with severe acute respiratory syndrome. J Infect Dis 193, 792–795 (2006).

28. I. Hamming et al., Tissue distribution of ACE2 protein, the functional receptor for SARS coronavirus. A first step in understanding SARS pathogenesis. J Pathol 203, 631–637 (2004).

29. J. Gu et al., Multiple organ infection and the pathogenesis of SARS. J Exp Med 202, 415–424 (2005).

30. Y. Ding et al., Organ distribution of severe acute respiratory syndrome (SARS) associated coronavirus (SARS-CoV) in SARS patients: implications for pathogenesis and virus transmission pathways. J Pathol 203, 622–630 (2004).

31. V. D. Menachery et al., Trypsin treatment unlocks barrier for zoonotic bat coronaviruses infection. J Virol, (2019).

32. Y. He, J. Li, S. Heck, S. Lustigman, S. Jiang, Antigenic and immunogenic characterization of recombinant baculovirus-expressed severe acute respiratory syndrome coronavirus spike protein: implication for vaccine design. J Virol 80, 5757–5767 (2006).

33. J. T. Lin et al., Safety and immunogenicity from a phase I trial of inactivated severe acute respiratory syndrome coronavirus vaccine. Antivir Ther 12, 1107–1113 (2007).

## Supplementary references

1. C. Brinkmann et al., The glycoprotein of vesicular stomatitis virus promotes release of virus-like particles from tetherin-positive cells. PLoS One 12, e0189073 (2017).

2. M. Hoffmann et al., Differential sensitivity of bat cells to infection by enveloped RNA viruses: coronaviruses, paramyxoviruses, filoviruses, and influenza viruses. PLoS One 8, e72942 (2013).

3. H. Hofmann et al., Human coronavirus NL63 employs the severe acute respiratory syndrome coronavirus receptor for cellular entry. Proc Natl Acad Sci U S A 102, 7988-7993 (2005).

4. S. Bertram et al., TMPRSS2 and TMPRSS4 facilitate trypsin-independent spread of influenza virus in Caco-2 cells. J Virol 84, 10016–10025 (2010).

5. S. Gierer et al., The spike protein of the emerging betacoronavirus EMC uses a novel coronavirus receptor for entry, can be activated by TMPRSS2, and is targeted by neutralizing antibodies. J Virol 87, 5502–5511 (2013).

6. H. Kleine-Weber et al., Mutations in the Spike Protein of Middle East Respiratory Syndrome Coronavirus Transmitted in Korea Increase Resistance to Antibody-Mediated Neutralization. J Virol 93, (2019).

7. M. Berger Rentsch, G. Zimmer, A vesicular stomatitis virus replicon-based bioassay for the rapid and sensitive determination of multi-species type I interferon. PLoS One 6, e25858 (2011).

